# Rapid estimation of cortical neuron activation thresholds by transcranial magnetic stimulation using convolutional neural networks

**DOI:** 10.1101/2022.05.18.490331

**Authors:** Aman S. Aberra, Adrian Lopez, Warren M. Grill, Angel V. Peterchev

**Author notes:** Corresponding author: A.V. Peterchev, Address: Division of Brain Stimulation and Neurophysiology, Department of Psychiatry and Behavioral Sciences, Duke University, Box 3620, DUMC, Durham, NC 27710, USA. Email addresses, and.

## Abstract

**Background:** Transcranial magnetic stimulation (TMS) can modulate neural activity by evoking action potentials in subpopulations of cortical neurons. The TMS-induced electric field (E-field) can be simulated in subject-specific head models derived from MR images, but the spatial distribution of the E-field alone does not predict the physiological response. Coupling E-field models to populations of biophysically realistic neuron models yields insights into the activation mechanisms of TMS, but the significant computational cost associated with these models limits their use and eventual translation to clinically relevant applications.

**Objective:** The objective was to develop computationally efficient estimators of the activation thresholds of multi-compartmental cortical neuron models in response to TMS-induced E-field distributions.

**Methods:** Multi-scale models combining anatomically accurate finite element method (FEM) simulations of the TMS E-field with layer-specific representations of cortical neurons were used to generate a large dataset of activation thresholds. 3D convolutional neural networks (CNNs) were trained on these data to predict the activation threshold of specific model neurons given the local E-field distribution. Using training and test data from different head models, the CNN estimator was compared to an approach using the uniform E-field approximation to estimate thresholds in the non-uniform TMS-induced E-field.

**Results:** The 3D CNNs were more accurate than the uniform E-field approach, with mean absolute percent error (MAPE) on the test dataset below 2.5% compared to 5.9 – 9.8% with the uniform E-field approach. Further, there was a strong correlation between the CNN predicted and actual thresholds for all cell types (*R*^2^ > 0.96) compared to the uniform E-field approach (*R*^2^ = 0.62 – 0.91). The CNNs estimate thresholds with a 2 – 4 orders of magnitude reduction in the computational cost of the multi-compartmental neuron models.

**Conclusion:** 3D CNNs can estimate rapidly and accurately the TMS activation thresholds of biophysically realistic neuron models using sparse samples of the local E-field, enabling simulating responses of large neuron populations or parameter space exploration on a personal computer.

## 1. Introduction

Transcranial magnetic stimulation (TMS) is a technique for noninvasive modulation of brain activity in which an electric field (E-field) is induced in the head by a current pulse applied through an external coil [1]. TMS is FDA-cleared to treat depression, obsessive compulsive disorder, smoking addiction, and migraine [2–7], and is under investigation for numerous other psychiatric and neurological disorders [8]. In addition, non-invasive stimulation of the cerebral cortex with TMS is valuable for human neuroscience research [9]. Nonetheless, TMS suffers from several limitations including large inter- and intra-subject variability in responses [10,11] and modest effect sizes relative to those achieved, for example, by electroconvulsive therapy [12]. Improving TMS efficacy and reliability is difficult using empirical methods alone due to the vast parameter space and limited understanding of the neural mechanisms by which TMS activates neurons and produces long-lasting changes to excitability.

Computational modeling of the TMS-induced E-field distribution in subject-specific volume conductor head models enables quantifying the E-field delivery to cortical targets [13–17]. However, the E-field distribution alone does not predict the response to stimulation, particularly when considering temporal characteristics of the stimulus (e.g., pulse shape and direction) or the diversity of neural elements in the brain. How TMS affects neural activity is still unclear and presents a complex problem, as the cortex is composed of various cell types, differing in morphology, electrophysiology, and connectivity, which all can contribute to the responses evoked by stimulation.

Previously, we developed models of human cortical neurons to study the response to the simulated TMS E-field [18,19]. This multi-scale modeling framework computes the polarization and activation of neural elements in response to arbitrary coil geometries and placements, as well as pulse waveforms and successfully reproduced trends in TMS thresholds as a function of pulse shape, width, and direction [18]. However, this approach has considerable computational cost of solving numerically the large system of partial differential equations associated with each neuron model, and requires high performance computing (HPC) resources when simulating large populations of neurons or variations in stimulus parameters. For example, simulating the threshold of a single neuron in our model [18] required 5–15 seconds; therefore, simulating the response of all neurons in the precentral gyrus (approximately 154 million^1^) to a TMS pulse would require 24–73 years run serially on a typical laptop or 3–10 months if parallelized over 100 CPUs. Thus, alternative approaches are required to advance models of the effects of TMS on neurons.

Machine learning provides a potential alternative to generate accurate, computationally efficient estimators of the neural response. Artificial neural networks (ANNs) and deep learning have achieved substantial success in multiple problem domains involving complex, high-dimensional data [23]. Convolutional neural networks (CNNs) are a class of deep, feed-forward ANNs that use convolutional kernels to extract local features from spatially structured data [23]. 3D CNNs were used to estimate TMS-induced E-field distributions in real-time [24,25] and to learn the mapping between the E-field distribution and evoked muscle responses [26]. CNNs were also used to learn the input–output properties of single neuron models for synaptic inputs [27,28], but they have yet to be applied to estimating neural activation by extracellular E-fields.

We designed a CNN that learned the mapping between TMS induced E-field distributions and the firing responses of biophysically realistic, multicompartmental model neurons, providing a rapid, computationally efficient method to quantify neural activation within E-field volume conductor models. We evaluated the performance of the CNNs and found them to produce accurate estimates of activation threshold, with mean absolute percent error close to the 2% window used in the simulated threshold binary search, in comparison to a simpler estimation approach using the uniform E-field approximation, which had mean absolute percent error of over 6%. Crucially, the CNN estimators ran 2–4 orders of magnitude faster than the full neuronal simulations.

## 2. Methods

We developed computationally efficient CNN estimators of the activation thresholds of biophysically realistic, multi-compartmental cortical neuron models in response to a TMS-induced electric field in subject-specific FEM head models derived from MRI data. After determining activation thresholds in the biophysically realistic neuron models, we trained a CNN to take as input the local E-field at regularly defined points around a neuron and output the threshold E-field magnitude to activate the neuron. CNNs were trained on thresholds for neurons placed in one head model (*almi5*) and tested on neuron thresholds from another head model (*ernie*).

### 2.1. Multi-scale model of TMS-induced cortical activation

The “ground truth” neural responses were obtained by simulating biophysically realistic models of L2/3 pyramidal cells (PCs), L4 large basket cells (LBCs), and L5 PCs coupled to E-fields computed within two MRI-derived volume conductor head models in SimNIBS v3.1 [29].

#### 2.1.1. E-field model

Two tetrahedral FEM meshes were generated using the *almi5* and *ernie* datasets included with SimNIBS, consisting of both T1- and T2-weighted images and diffusion tensor imaging (DTI) data. We used the *mri2mesh* pipeline [30] with white matter surface resolution set to 60,000 vertices for the *almi5* dataset and 120,000 vertices for the *ernie* dataset. The *almi5* volume head mesh consisted of 646,359 vertices and 3.6 million tetrahedral elements, and the *ernie* volume head mesh consisted of 1.6 million vertices and 8.8 million tetrahedral elements. The meshes consisted of five homogenous compartments: white matter, gray matter, cerebrospinal fluid (CSF), bone, and scalp. For the *ernie* mesh, all compartments were assigned default conductivity values, with anisotropic conductivity in the white matter using the DTI data and the volume normalized approach (mean conductivity = 0.126 S/m), and isotropic conductivities in the other tissues: gray matter: 0.275 S/m, CSF: 1.654 S/m, bone: 0.01 S/m, scalp: 0.25 S/m. All conductivities were the same in the *almi5* mesh, except for the gray matter (0.276 S/m) and CSF (1.79 S/m), which were the values used in our previous publication [18].

E-field distributions were simulated for the MC-B70 figure-of-8 coil (P/N 9016E056, MagVenture A/S, Farum, Denmark), which has ten turns in each of the two windings with outer and inner diameters of 10.8 and 2.4 cm, respectively [31]. The coil was positioned in both cases above the left motor hand knob, located on the precentral gyrus [32]. For the *almi5* mesh, we simulated both posterior–anterior (P–A) and latero–medial (L–M) coil orientations, with the coil handle oriented 45° and 90° relative to the midline, respectively. Using these two orientations, the E-field distributions for the A–P and M–L pulse directions were generated by flipping all E-field vectors. For the *ernie* mesh, we simulated the P–A coil orientation of the motor hand knob. The E-field distributions were computed with a coil-to-scalp distance of 2 mm and coil current of 1 A/µs.

#### 2.1.2. Neuron models

Previously, we adapted the multi-compartmental, conductance-based models of juvenile (P14) rat cortical neurons implemented by the Blue Brain Project [33,34] to the biophysical and geometric properties of adult, human cortical neurons, and implemented them in NEURON [35]. TMS activated with lowest intensity the L5 and L2/3 PCs as well as L4 large basket cells (LBCs); therefore, the current study focused on these neurons (we focused our simulations on the L4 LBCs, but preliminary simulations indicated LBCs in other layers had similar thresholds [18]). Each cell type had five “virtual clones”, which had stochastically varied morphologies but identical biophysical parameters [18,19]. The cell morphologies are plotted in Supplementary Figure S1.

#### 2.1.3. Embedding neuron populations in head model

Regions of interest (ROIs) were defined within each head mesh to embed layer-specific populations of model neurons. The *almi5* mesh ROI was defined as a 32 × 34 × 50 mm^3^ region containing the M1 hand knob on the precentral gyrus and opposing postcentral gyrus [18]. For the *ernie* mesh, a larger ROI was defined to include the precentral gyrus, central sulcus, and postcentral gyrus labeled regions generated by Freesurfer’s automatic cortical parcellation with the Destrieux Atlas [36]. These regions were cropped with a 48 × 55 × 50 mm^3^ box. Neuron models were positioned and oriented within the gray matter by interpolating surface meshes representing each cortical layer between the gray matter and white matter surfaces and discretizing the surfaces with the number of triangular elements matching the desired number of neurons in each layer. For the *almi5* and *ernie* ROIs, 3,000 and 5,000 elements were used for each layer, resulting in surfaces with mean density of 1.7 and 0.64 elements (i.e., neuron positions) per mm^2^, respectively. The layer depths were defined using layer depth boundaries from the recently published layer segmentation of the BigBrain histological atlas; in von Economo area FA, the boundaries between adjacent layers were at normalized depths of 0.0993 (L1–L2/3), 0.466 (L2/3–L4), 0.524 (L4–L5), 0.753 (L5–L6) (total depth of gray matter is 1) [37]. Accordingly, the cell placement surface meshes were positioned between these boundaries at normalized depths of L2/3: 0.4, L4: 0.5, L5: 0.75 for the *ernie* model and L2/3: 0.4, L4: 0.55, L5 0.65 for the *almi5* model. The slight differences in the tissue conductivity values and layer depths add to the anatomical variation between the two head models which was advantageous for testing of the robustness of the CNN estimators.

Single model neurons were placed with their cell bodies centered in each element and oriented to align their somatodendritic axis normal to the element. Using the somatodendritic axis as the polar axis of the local spherical coordinate system (Figure 1D), each model neuron was placed with initial random azimuthal orientation and then rotated to 11 additional orientations with 30° steps to sample the full range of possible orientations and generate a larger dataset for training and evaluating the CNNs (discussed in Section 2.2.4).

**Figure 1.**
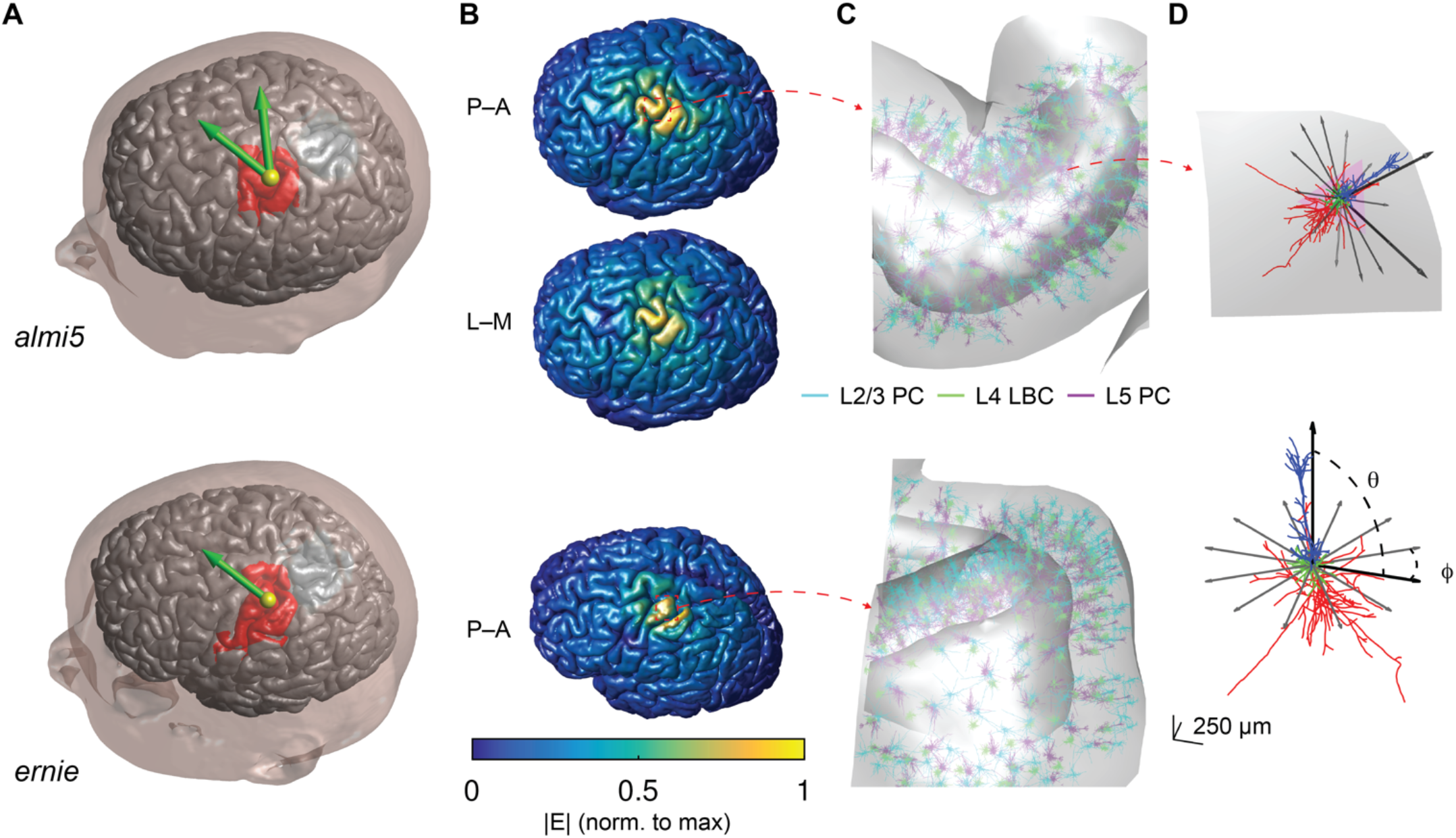
Multi-scale model of TMS-induced activation. **A)** FEM head models used in this study to compute E-fields with simulated TMS coil positions (yellow sphere is location of coil center) and directions (green arrow points opposite of coil handle). Model neurons were populated throughout ROI (red region) encompassing the motor hand knob and opposing postcentral gyrus. **B)** E-field magnitudes plotted on gray matter surface for P–A and L–M coil orientations (top) and P–A (bottom) in corresponding meshes. **C)** Neuron populations in corresponding head meshes shown with zoomed in view. **D)** Example L5 PC placed in gyral crown (top), oriented with the somatodendritic axis normal to the element and reference vectors indicating azimuthal rotations simulated at all positions (tangential to element normal). Same L5 PC model neuron shown in cell-centered coordinate system with somatodendritic axis aligned to polar axis (bottom) with polar angle *θ* and azimuthal angle *ϕ*. The azimuthal rotations shown are the same as in the top panel.

Mesh generation, placement of neuronal morphologies, extraction of E-field vectors from the SimNIBS output, NEURON simulation control, analysis, and visualization were conducted in MATLAB (R2016a & R2017a, The Mathworks, Inc., Natick, MA, USA).

#### 2.1.4. Neuron simulations

Applying the quasi-static approximation [38,39] allows the separation of the spatial and temporal components of the TMS-induced E-field. The spatial component was derived from the E-field distributions computed in SimNIBS with a coil current rate of change of 1 A/µs by interpolating the E-field vectors at each model neuron’s compartments after placement within the head mesh. The E-field at the model neuron compartments was linearly interpolated from the 10 nearest mesh points (tetrahedral vertices in SimNIBS) within the gray and white matter volumes using the MATLAB `scatteredInterpolant` function. The E-field vectors were integrated along each neural process to generate a quasipotential [40–42], which was coupled as an extracellular voltage to each compartment in NEURON using the `extracellular` mechanism [35]. The neuron models were discretized with isopotential compartments no longer than 20 µm.

The temporal component of the E-field was included by scaling uniformly the quasipotentials over time by either a monophasic or biphasic TMS pulse recorded from a MagPro X100 stimulator (MagVenture A/S, Denmark) with a MagVenture MCF-B70 figure-of-8 coil (P/N 9016E0564) using a search coil and sampling rate of 5 MHz. The E-fields were down-sampled to twice the simulation time step and normalized to unity amplitude for subsequent scaling in the neural simulations. We used a simulation time step of 5 µs, simulation window of 1 ms, and backward Euler integration. Activation thresholds were determined by scaling the pulse waveform using a binary search algorithm to find the minimum stimulus intensity, within 2%, necessary to elicit an action potential, defined as the membrane potential in at least 3 compartments crossing 0 mV with positive slope.

### 2.2. Convolutional neural network for threshold estimation

We used a two-stage 3D CNN followed by 1D dense layers to estimate the threshold to activate each model neuron (Figure 2). The CNN does not represent the temporal dynamics of the neural response; therefore, each CNN estimates the activation thresholds for the specific TMS pulse waveform that was used to generate the training data. CNNs were trained on thresholds for neurons placed in the *almi5* model and tested on neuron thresholds from the *ernie* model. Hyperparameters were tuned using random search with the training dataset. The result was a set of 15 trained CNNs for each of the 15 model neurons, with each CNN outputting the activation threshold of a model neuron for any local E-field distribution and the specific pulse waveform.

**Figure 2.**
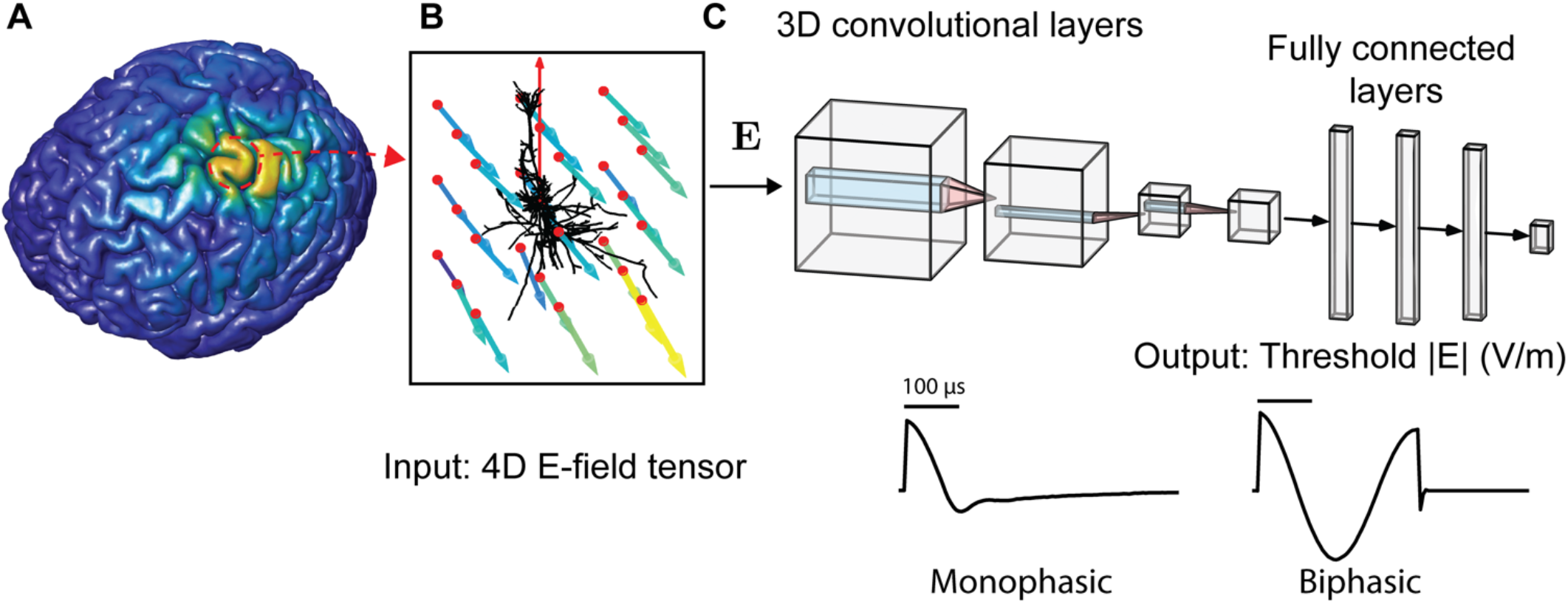
Estimating E-field threshold of multi-compartmental neurons using convolutional neural networks (CNNs). **A)** E-field distribution computed throughout head model using SimNIBS, shown on gray matter surface. **B)** For a model neuron at any location, local E-field vectors were sampled at ***N*** × ***N*** × ***N*** regular grid (red points) centered on cell body and rotated into cell-centered coordinate system (red vector indicates somatodendritic axis, i.e., polar axis). Color and length of each vector indicates magnitude. **C)** After normalizing the E-field vector magnitudes by that of the grid central node, the E-field vectors were structured as a 4D tensor (*N* × *N* × *N* × 3) input to the 3D convolutional layers. The output of the final 3D convolutional layer was flattened and input into dense, fully connected layers, which ended with a single linear output layer element with the predicted threshold E-field magnitude in V/m. E-field magnitude is referenced to E-field at the central grid point, as off-center points have, in general, different direction/magnitude. Separate models were trained on thresholds obtained with different pulse shapes, either a monophasic or biphasic TMS pulse, shown below.

#### 2.2.1. E-field input and preprocessing

The input to the CNN was E-field vectors defined on an *N* × *N* × *N* cubic grid with side length *l* centered on the cell body and rotated into the cell-centered coordinate system, comprising a 4D tensor (*N* × *N* × *N* × 3). This ensured the spatial relationship between the E-field sampling points and the morphology was constant for any model neuron placed in the brain. We set *l* to encompass approximately the cell dimensions while minimizing grid points penetrating the CSF: 2 mm for L2/3 PCs, 1.5 mm for L4 LBCs, and 1.5 mm for L5 PCs. The effect of varying the number of grid points and grid size was tested with the L5 PCs, using *N* = 3, 5, 7, and 9 points per dimension and *l* = 1 to 3 mm in 0.5 mm steps. The sampling grids used in the main results for all cells are shown in Supplementary Figure S1.

The E-field tensors were input to the CNN with E-field components in either Cartesian coordinates (*E*_*x*_, *E*_*y*_, *E*_*z*_) or spherical coordinates (*E*_*r*_, *E*_*θ*_, *E*_*ϕ*_), again using the somatodendritic axis as the polar axis of the local spherical coordinate system. Since the azimuthal angle ranged from 0–360°, creating a periodic discontinuity at 0°, the azimuthal angle was separated into two components ranging from −1 to 1 using cos and sin, i.e., (*E*_*r*_, *E*_*θ*_, cos *E*_*ϕ*_, sin *E*_*ϕ*_). We hypothesized using spherical coordinates would enable the CNN learn the strong dependence of model thresholds on the E-field magnitude [18], which is included explicitly in the spherical coordinate representation.

#### 2.2.2. Local E-field characterization

To determine how well the E-field sampling grids represented spatial gradients of the E-field from the FEM solution, we computed the magnitude of the directional gradients at each neuronal position, given by the norm of the E-field Jacobian,

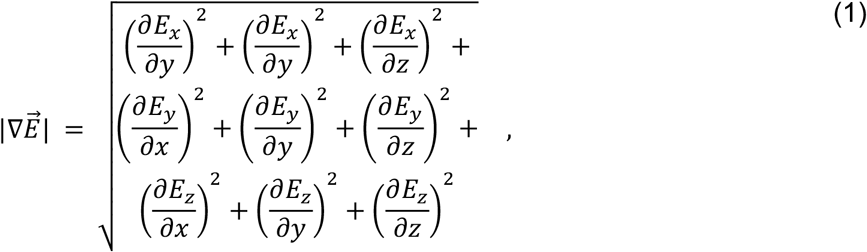

and extracted the median value of this metric within each sampling grid size *N*. The E-field vectors at each grid point were first divided by the magnitude of the E-field at the center grid point (cell body). We also computed this gradient metric with an *N* = 13 sampling grid to capture better the higher spatial frequencies in the FEM solution. Additionally, we quantified how well the E-field sampling grids captured the E-field at the action potential (AP) initiation site by linearly interpolating the E-field vector within each sampling grid at the AP initiation site, extracted from the NEURON simulations. We compared this interpolated E-field vector 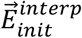 to the actual E-field vector from the FEM solution 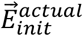 by computing the absolute percent error in magnitude 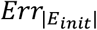 and error in angle *θ*_*err,init*_ between them, given by

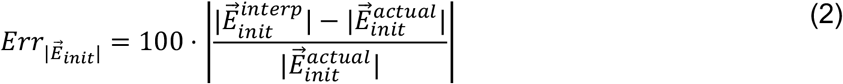

and

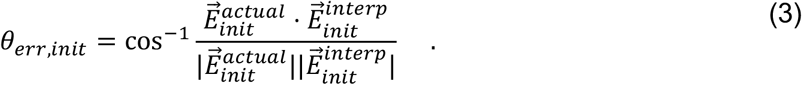

#### 2.2.3. CNN architecture

The architecture consisted of a series of 3D convolutional layers (Conv3D) followed by a flattening operation (Flatten layer) and a series of fully connected, 1D dense layers (Dense) and a final one-unit output layer corresponding to the estimated threshold E-field magnitude. This threshold can then be converted to the threshold TMS device intensity setting by scaling with the ratio between the coil current rate of change and the local E-field magnitude from the FEM simulation. The model weights (both convolutional kernels and dense layers) were initialized with Xavier uniform initialization [43] (glorot_uniform) and trained using the Adam adaptive, stochastic gradient-based optimization algorithm [44] with the mean squared error loss function. The CNNs were implemented using Tensorflow 2.2 [45] with the Keras deep-learning API in Python 3.7.

ANN hyperparameters are still commonly tuned by hand based on intuition, due to the vast parameter space and often slow training times required to evaluate each selected hyperparameter set. Here, we tuned the hyperparameters of the general architecture described above (e.g., number of convolutional layers, dense layers, etc.) using random search, based on previous work that it was more efficient than grid search [46], followed by some manual tuning of individual hyperparameters. Each set of hyperparameters tested in the random search was used to train a candidate CNN architecture on a third of the training dataset to speed up training time. The search ranges and best hyperparameters obtained are included in

. Using the `RandomizedSearchCV` function in the scikit-learn Python module [47], a total of 52 hyperparameter sets were randomly selected and evaluated based on their final test dataset loss (MSE) after 1000 epochs for each of the L5 PC clones, without using cross validation to reduce search time. We first conducted the hyperparameter search for the five L5 PCs and found that model performance was always the same across clones. Therefore, we ran random searches for a single L2/3 PC and L4 LBC clone and found that the best hyperparameters from the first search with the L5 PCs were not outperformed by any new hyperparameter sets tested.

After exploring the hyperparameter space, we found the best performing CNN architecture for the 9 × 9 × 9 E-field sampling grid consisted of four convolutional layers and three dense layers with 3 × 3 × 3 convolutional kernels (Table 1). The number of filters and dense units were defined for the initial convolutional and dense layer, respectively, and then decreased in each subsequent layer by multiplying the first layer’s value (115 filters and 57 dense units) by a shrink rate *r*^*i*−1^, where *i* is the index of the convolutional or dense layer. The output of each convolutional and dense layer was passed through rectified linear units (`ReLU`). The convolutional layers all had kernels of size 3 × 3 × 3 (stride of 1) and did not use zero-padding, resulting in layers with decreasing dimension: for the 9 × 9 × 9 E-field vector sampling grid, the first three dimensions of the layer outputs were 7, 5, 3, and 1. For the models using fewer E-field sampling points (*N* = 3, 5, 7), the dimensions of the input were incompatible with this architecture, so we used two approaches to adapt the network to these inputs. The first was to reduce the kernel size to 2 × 2 × 2, and, for the *N* = 3 case, reduce the number of convolutional layers to two (variable network size). However, since this changed the overall number of weights, we tested a second approach in which the architecture was kept constant and the E-fields were upsampled to the 9 × 9 × 9 grid using linear interpolation (constant network size).

**Table 1.** Convolutional neural network (CNN) hyperparameters. Columns include tuning range randomly searched for each hyperparameter and best hyperparameters identified.

| Hyperparameter | Tuning range | Best |
| --- | --- | --- |
| Number of convolutional layers | 2 – 6 | 4 |
| Convolutional kernel size | 1 – 3 | 3 |
| Number of convolutional filters (1<sup>st</sup> layer) | 50–100 | 115 |
| Number of dense layers | 1–5 | 3 |
| Number of dense units (1<sup>st</sup> layer) | 50 – 1000 | 57 |
| Shrink rate | 0.5 – 0.9 | 0.8 |
| Batch size | 8–128 | 62 |
| Learning rate (initial) | $10^{-8} - 10^{-5}$ | $10^{-5}$ |
| Activation function | ReLU or leaky ReLU | ReLU |
| Dropout rate | 0 – 0.5 | 0 |
| Batch normalization | off or on | off |
| Output activation function | – | linear |

Training was conducted using mini-batches of 62 samples and an initial learning rate of 10^−5^. To train the final set of CNNs with the best parameter set, we used a maximum of 2000 epochs, and training was terminated to reduce overfitting if the validation loss did not decrease after 30 epochs (`EarlyStop`). We also used learning rate scheduling to reduce the learning rate by a factor of 5 if the validation loss plateaued for 15 epochs (`ReduceLRonPlateau`). Example training curves are shown in Supplementary Figure S2.

We tested including dropout layers between each dense layer, as well as batch normalization either before or after the non-linearity for regularization. These operations did reduce training time, but they resulted in higher prediction error and were not included in the final hyperparameter set. The same hyperparameter set was then used for each of the 15 CNNs corresponding to the model neurons.

#### 2.2.4. Training and testing datasets

The training and test datasets for each model neuron consisted of pairings of 4D E-field tensors sampled around each model neuron position within the cortical geometry and the corresponding activation thresholds. For each of the 11 additional azimuthal rotations, the E-field sampling grids were rotated to ensure the E-field vector orientations relative to the model neuron were constant, while providing a unique E-field distribution for training the CNN. The training dataset was derived from the E-field simulations in the *almi5* mesh, consisting of four stimulation directions (P–A, A–P, L–M, M–L), 2,999–3,000 neuron positions per cell, and 12 rotations, totaling 143,952 or 144,000 unique E-field– threshold combinations for each of the 15 model neurons (2.16 million simulations). This training dataset was split, with 85% of the data used for training the model and 15% used for validation. CNNs were trained on HPC cluster nodes each equipped with a GeForce RTX 2080 Ti graphics card.

To test model generalization to a new head model, the test dataset consisted of E-field distributions from two stimulation directions (P–A and A–P) in the *ernie* mesh, with 4,999 neuron positions, and 12 rotations at each position, totaling 119,976 unique E-field–threshold combinations for each model neuron. After training with the training/validation dataset, the performance of each CNN was evaluated on this test dataset.

### 2.3. Method for estimating TMS activation thresholds with uniform E-field simulations

We implemented an additional method to estimate the neuron model thresholds that approximated the local E-field as uniform. The thresholds were pre-simulated for uniform E-field applied at range of directions spanning the polar and azimuthal directions. To simulate the response to uniform E-field, the extracellular potential *V*_e_ was computed at each compartment with position (*x, y, z*) using

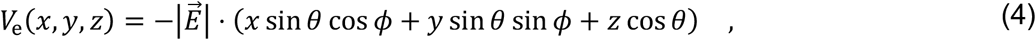

where the direction of the uniform E-field was given by polar angle *θ* and azimuthal angle *ϕ*, in spherical coordinates with respect to the somatodendritic axes (Figure 1D), and the potential of the origin (soma) was set to zero, as in [18,19]. Uniform E-field was applied with the monophasic or biphasic TMS pulse at each direction with steps of 5° for both the polar and azimuthal directions, for a total of 2,522 directions, generating threshold–direction maps for each model neuron.

The threshold–direction maps were then used to estimate thresholds of neurons embedded in the non-uniform FEM E-field. For a given neuron, the E-field vector at the soma was extracted and rotated into the cell-centered coordinate system used in the uniform E-field simulations (Figure 1D). The threshold (in V/m) for the corresponding E-field direction was then interpolated from the uniform E-field threshold–direction map. This threshold was divided by the FEM E-field vector’s magnitude per A/µs current rate of change to convert to stimulator output intensity. This latter step was only necessary to determine thresholds in terms of stimulator intensity, which would be the relevant “knob” for an experimenter, rather than local E-field intensity. The threshold–direction map interpolant was implemented in MATLAB as a `griddedInterpolant` using first-order (linear) interpolation.

### 2.4. Code and data availability

The code and relevant data of this study will be made available on GitHub upon publication.

## 3. Results

### 3.1. CNN predicts accurately the threshold response of single neurons to TMS

The best CNN architecture provided remarkably accurate predictions of the activation thresholds for L2/3 PC, L4 LBC, and L5 PC neurons. Figure 3A shows the spatial distribution of thresholds for monophasic, P–A TMS of M1 in the *ernie* head model (test dataset) calculated in NEURON, predicted with the ML trained on the *almi5* head model, and estimated with the uniform E-field approximation. The relative errors were substantially lower for the CNN compared to the uniform E-field method (Figure 3B). Median errors across clones and rotations for the L4 LBCs and L5 PCs ranged from −3.4 to 6.6% and −1.9% to 3.5%, respectively, for the CNN, and −29.5% to 38.8% and −27.5 to 49.5%, respectively, for the uniform E-field method. The L2/3 PCs had the largest errors for either approach, with median error ranging from −23.7% to 37.1% with the CNN and −31.9% to 97.1% with the uniform E-field method. These outer extrema of the median error distributions were driven by outliers, with 97.3% below 5% error for the CNN, compared to only 59.2% for the uniform E-field method. The full error distributions across layers for both methods are shown in Supplementary Figure S3.

**Figure 3.**
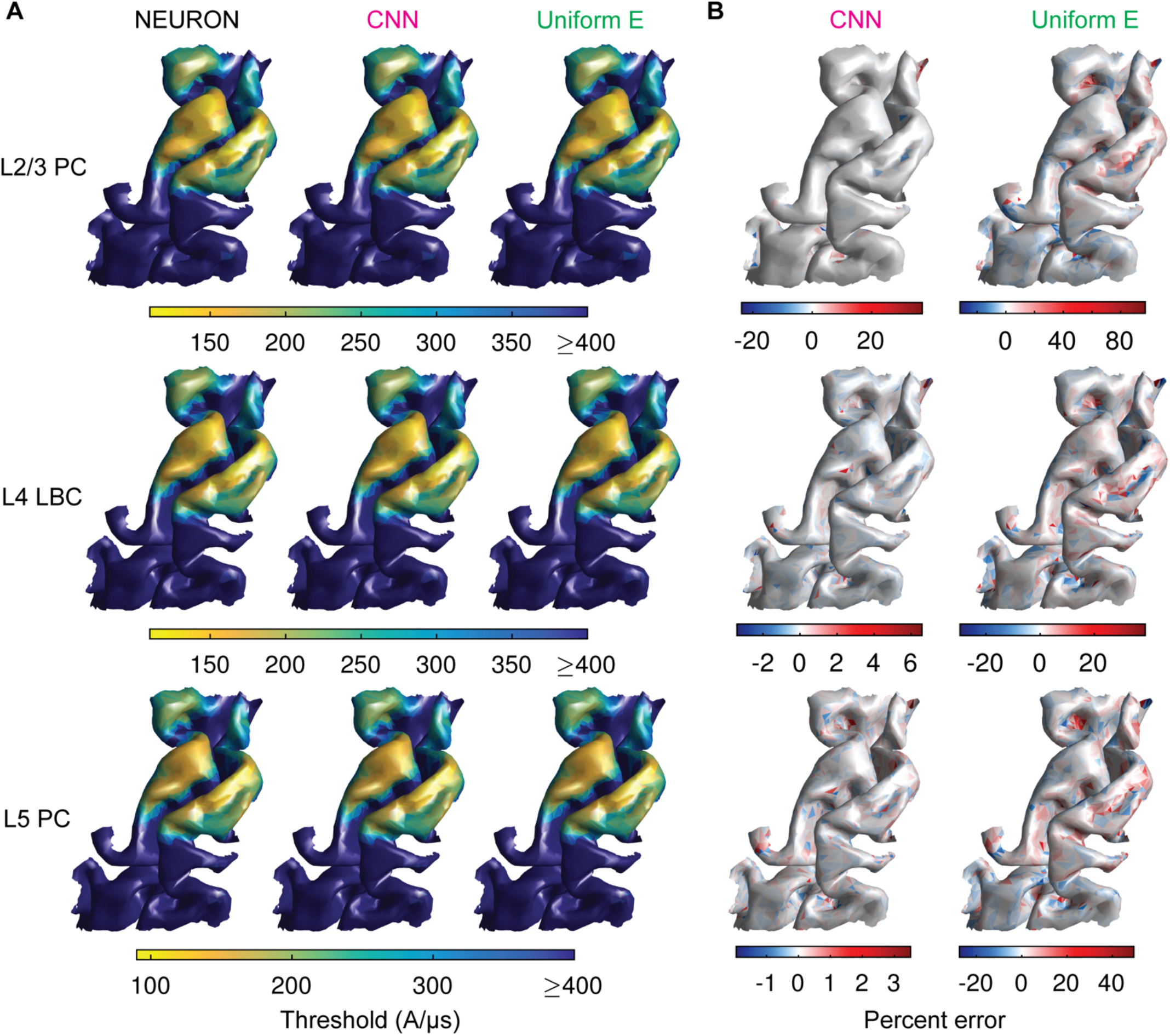
CNN accurately predicts thresholds for activation of neurons across the cortex. Actual and predicted thresholds for monophasic, P–A TMS of M1 in *ernie* head model (test dataset). A) Surface plots of median threshold stimulator intensity (in coil current rate of change) across clones and rotations of L2/3 PCs (top row), L4 LBCs (middle row), and L5 PCs (bottom row) for NEURON simulations (left column), CNN prediction (middle column), and uniform E-field method (right column). B) Surface plots of median percent error of thresholds across clones and rotations of L2/3 PCs (top row), L4 LBCs (middle row), and L5 PCs (bottom row) for CNN prediction (left column), and uniform E-field method (right column). Note the different color bar limits for the error distributions with CNN and uniform E-field method.

The CNN-predicted thresholds had high correlations with the thresholds calculated by repeatedly solving the non-linear cable equations across the entire set of neurons and positions, with *R*^2^ ranging 0.959 – 0.980, 0.974 – 0.993, and 0.988 – 0.996 across the L2/3 PC, L4 LBC, and L5 PC clones, respectively (Figure 4A). The correlations for thresholds predicted with the uniform E-field method were much weaker, with *R*^2^ ranging 0.6371 – 0.8405, 0.618 – 0.938, and 0.645 – 0.914 across the L2/3 PC, L4 LBC, and L5 PC clones, respectively (Figure 4A). The CNN outperformed the uniform E-field approach for all model neurons, yielding mean absolute percent error (MAPE) less than 2.2% (L2/3 PCs), 1.1% (L4 LBCs), and 1.4% (L5 PCs), and median absolute percent errors were less than 1.2% (L2/3 PCs), 0.75% (L4 LBCs), and 0.74% (L5 PCs). For the uniform E-field approach, MAPE was less than 9.0% (L2/3 PCs), 5.9% (L4 LBCs), and 9.8% (L5 PCs) (Figure 4B), and median absolute percent errors were less than 5.4% (L2/3 PCs), 3.3% (L4 LBCs), and 4.5% (L5 PCs). We also trained a set of CNNs on thresholds obtained with biphasic TMS pulses and found their performance was similar to that of the monophasic pulse CNN estimators (Supplementary Figure S4).

**Figure 4.**
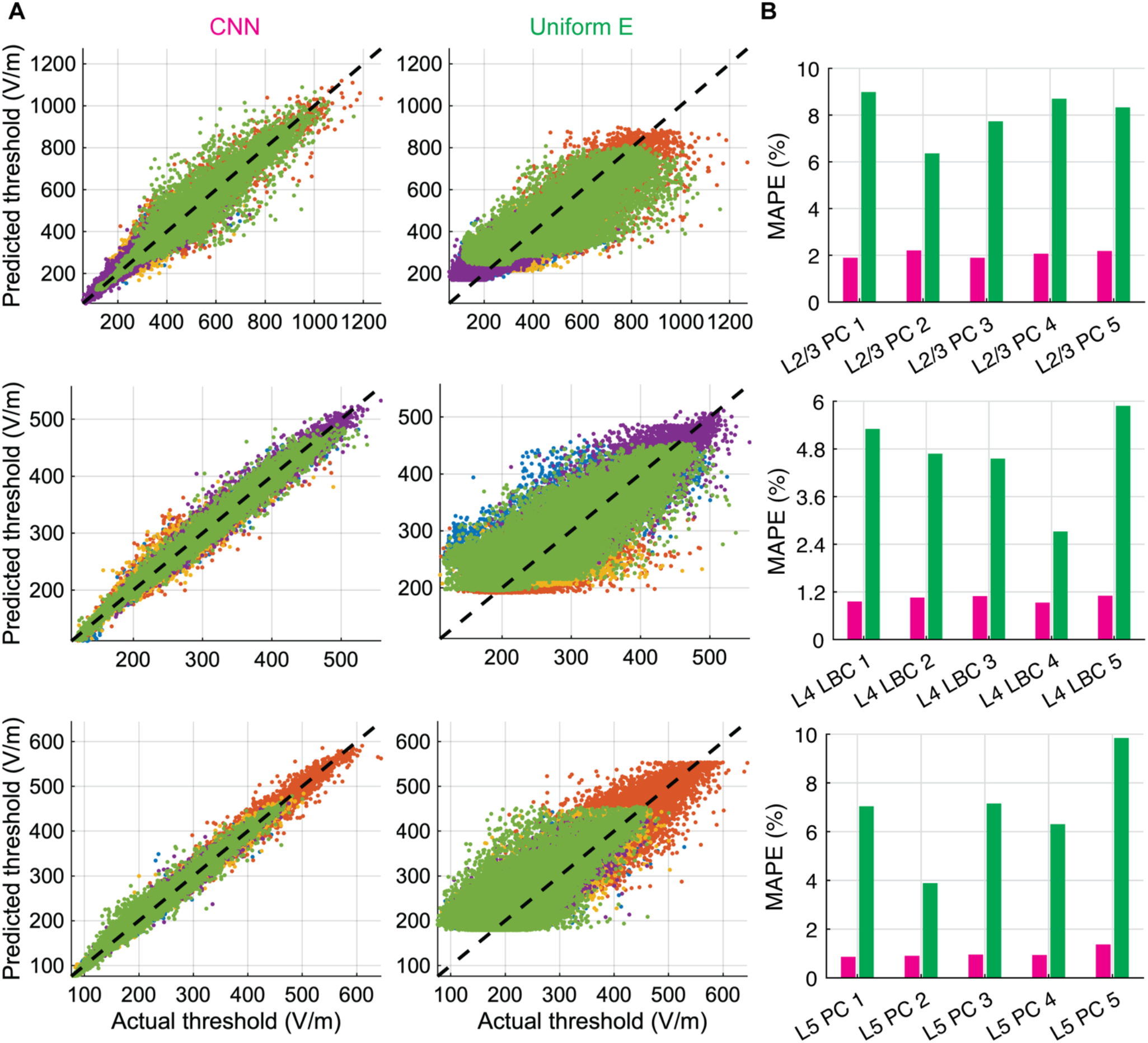
Distribution of threshold E-field errors for CNN and uniform E-field approximation. **A)** Predicted thresholds by CNN (left column) and uniform E-field approximation (middle column) across entire test dataset plotted against thresholds from NEURON simulations (actual) in magnitude of E-field at soma for L2/3 PCs (top row), L4 LBCs (middle row), and L5 PCs (bottom row). Different colors correspond to different clones within layer. **B)** Mean absolute percent error (MAPE) on test dataset for CNN (magenta) and uniform E-field approach (green), separated by clone within layer.

### 3.2. Dependence of CNN performance on E-field representation

The performance of ANNs often benefit from data pre-processing to assist in training or pre-identifying features that are known *a priori* to correlate with the output [48]; thus, we tested the effect of E-field representation and sampling parameters on CNN performance with the L5 PCs. Representing the E-field vectors in Cartesian coordinates yielded lower errors than spherical coordinates (Figure 5A). This held true for the L2/3 PCs and L4 LBCs as well (Supplementary Figure S5). Focusing on the L5 PCs, the optimal sampling grid size varied by clone, with lowest errors for the grid size that best matched the spatial extent of each specific axonal morphology (Figure 5B). The 1.5 mm grid produced the lowest mean error across all L5 PC clones, and we used this single grid size for the remaining results. Finally, reducing the number of sampling points increased the mean error, although MAPE was still below 4% for even the 3 × 3 × 3 sampling grid (27 E-field vectors) (Figure 5C). This trend held for both the variable and constant network sizes, with prediction errors differing by less than 1% (Supplementary Figure S6).

**Figure 5.**
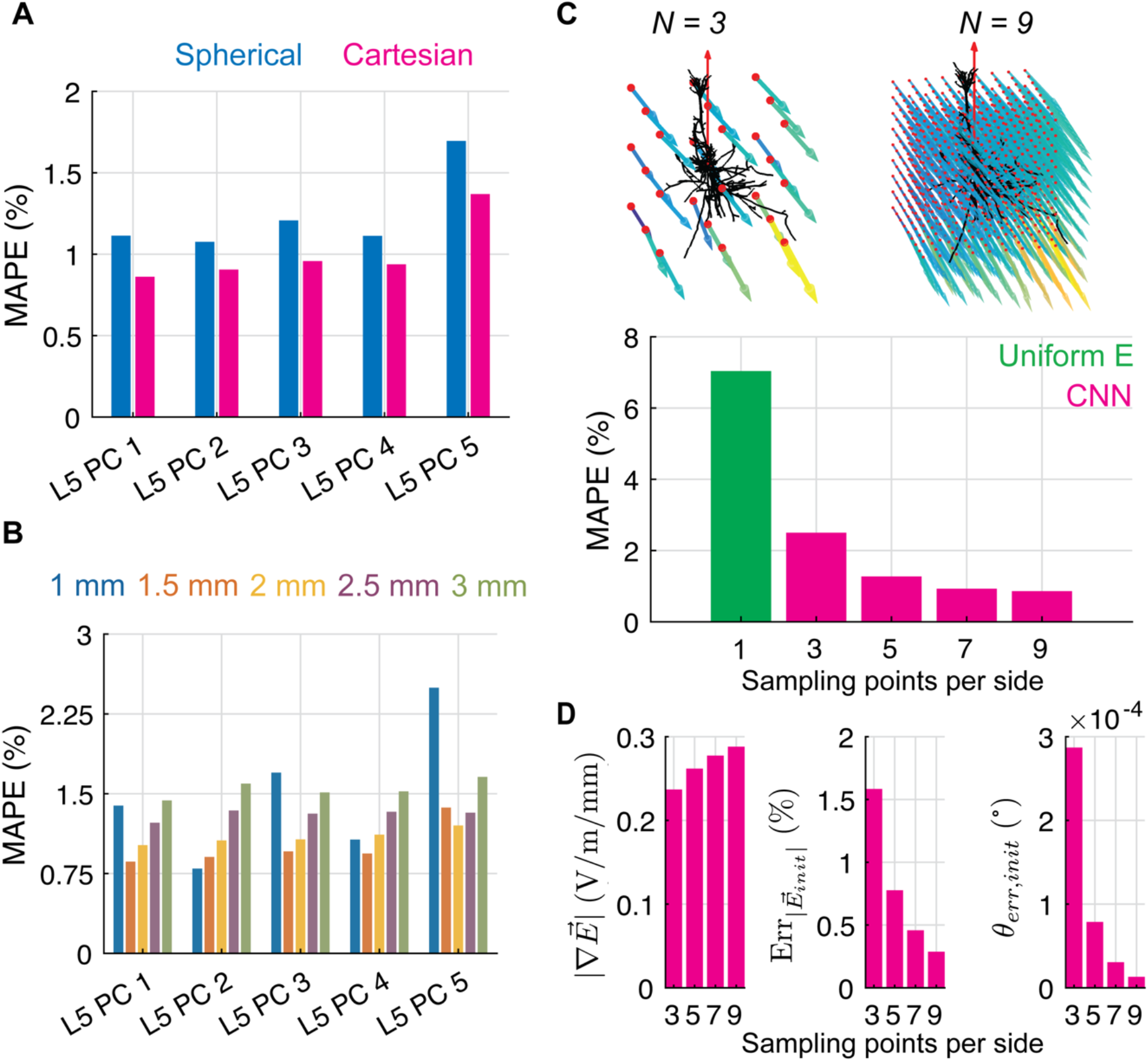
Effect of E-field representation on CNN performance. **A)** Mean absolute percent error metric on test dataset for L5 PC CNNs with E-field input in either spherical coordinates (represented with four variables; pre-processing described in Section 2.2.1) or Cartesian coordinates (represented with three variables). **B)** MAPE of test dataset for L5 PC CNNs for varying sampling grid sizes *l*. **C)** MAPE metric for example L5 PC (clone 1) CNN for different sampling resolutions *N* (magenta bars), using constant network size, with uniform E-field error metric included for comparison (green bar). **D)** Median magnitude of directional E-field gradients (left) and error in magnitude (middle) and direction (right) of the interpolated E-field at the AP initiation site in test dataset for different sampling resolutions *N* for L5 PC (clone 1).

We hypothesized that increasing sampling resolution improved CNN performance by more accurately representing local E-field gradients and the E-field magnitude and direction at the site of AP initiation. As expected, the FEM E-field was represented on average with lower spatial gradients when using fewer sampling points (Figure 5D, left), indicating higher spatial gradients of the E-field were lost due to undersampling. Furthermore, at the AP initiation site identified from the NEURON simulations, the E-field amplitude and direction were estimated less accurately using linear interpolation with fewer sampling points (Figure 5D, middle and right). We tested the contribution of these E-field metrics to prediction errors across the test dataset for each sampling resolution using linear regression (Supplementary Figure S7). Prediction errors were significantly correlated with all three metrics individually, with *R*^2^ = 0.14 for the median magnitude of the directional E-field gradients, *R*^2^ = 0.25 for the absolute percent error in the E-field magnitude at the AP initiation site, and *R*^2^ = 0.10 for the angle error of the E-field at the AP initiation site (all *N* = 3, *p* < 10^−5^) (Supplementary Figure S7D). Correlation strength was inversely related to sampling resolution, and multiple linear regression with all three metrics yielded *R*^2^ from 0.30 to 0.20 for *N* = 3 to 9, respectively. Of these metrics, the strongest contributor to the overall prediction error was the error in the interpolated E-field magnitude at the AP initiation site (*β* = 1.09, *N* = 3), followed by the E-field gradients (*β* = 0.58, *N* = 3) and error in the E-field direction at the AP initiation site (*β* = 0.11, *N* = 3) (Supplementary Figure S7E).

The performance on the test dataset demonstrated that the CNN could generalize to TMS thresholds for E-fields simulated in a head model not used in training, but the space of possible E-field distributions experienced by the model neurons may not be substantially different between head models. Therefore, we tested whether the CNNs could also predict response thresholds to uniform E-fields, which are essentially zeroth order polynomial components of the non-uniform E-fields seen in the training and test datasets. Figure 6A shows the threshold–direction maps for an example L5 PC generated with NEURON simulations and with the CNN; the CNN produced extremely low error, ranging from −2.5 to 1.9% across directions (Figure 6B).

**Figure 6.**
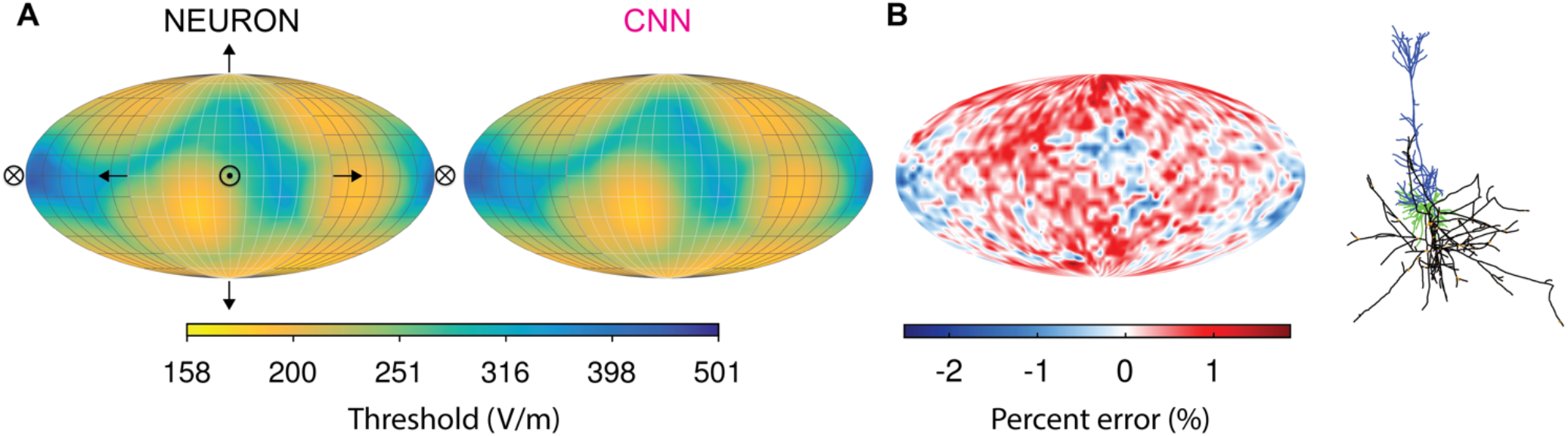
CNN trained on non-uniform TMS-induced E-field reproduces response to uniform E-field. **A)** Threshold–direction maps generated with NEURON simulations (left) and CNN estimation (right) for an example L5 PC. Mollweide projection of thresholds on a sphere in which normal vectors represent uniform E-field direction. Arrows indicate direction of E-field relative to cell (shown on right). Crossed circle represents E-field pointing into the page, and circle with dot represents E-field pointing out of the page. **B)** Percent error of CNN prediction. Inset: morphology of L5 PC used in this figure (blue are apical dendrites, green are basal dendrites, and black is axon).

The CNN trained on the TMS data also approximated the thresholds for a point current source (e.g., for intracortical microsimulation), reproducing qualitatively the current–distance relationship (Supplementary Figure S8). However, as expected, the prediction error was much higher than for TMS, as the point source generates a highly non-uniform E-field distribution. For comparison, the mean peak magnitude of directional gradients for the point source E-field was 482.3 V/m/mm, about 340 times higher than the mean peak gradient of the TMS-induced E-fields sampled with the 9 × 9 × 9 grid. Lower distances between the point source and activated neuronal compartment, which subject the neuron to higher spatial gradients, also led to higher errors (Supplementary Figure S8D).

### 3.3. CNN estimation provides massive speed up over NEURON simulations

We quantified the computational savings resulting from using the CNN to estimate thresholds compared to running simulations in NEURON on a single CPU of a typical laptop (Macbook Pro with 2.2 GHz i7-4770 CPU) or on an HPC machine with 76 CPUs (2.2 GHz Xeon Gold 6148) (Figure 7). Determining a single threshold with a binary search algorithm requires several evaluations, depending on the proximity of the initial guess to the threshold value; on a single CPU, this required 5.0 ± 0.2 sec for the L2/3 PCs (mean ± STD), 8.8 ± 0.5 sec for the L4 LBCs, and 9.5 ± 1.3 sec for the L5 PCs (n = 5 for all). In contrast, estimating thresholds for 1000 E-field inputs with the CNNs required 0.75 ± 0.17 ms per threshold (n = 5), which was similar between cell models due to the identical model architecture. Figure 7 illustrates how total time scales with the number of simulations (i.e., thresholds), showing sublinear increase in total time until ∼100 simulations for the CNN on a single CPU laptop, at which point total time increases linearly. In addition, Figure 7 depicts the best-case scenario for NEURON simulations by extrapolating total time based on the minimum time per simulation for each cell type if run on a single CPU or run in parallel on a 76 CPU HPC node with no parallelization overhead. By this conservative estimate, the CNN provides 2 to 4.2 orders of magnitude in computational savings compared to the NEURON simulations run serially^2^ and 2.1 to 2.7 orders of magnitude in computational savings compared to the NEURON simulations run in parallel on 76 CPUs on an HPC machine.

**Figure 7.**
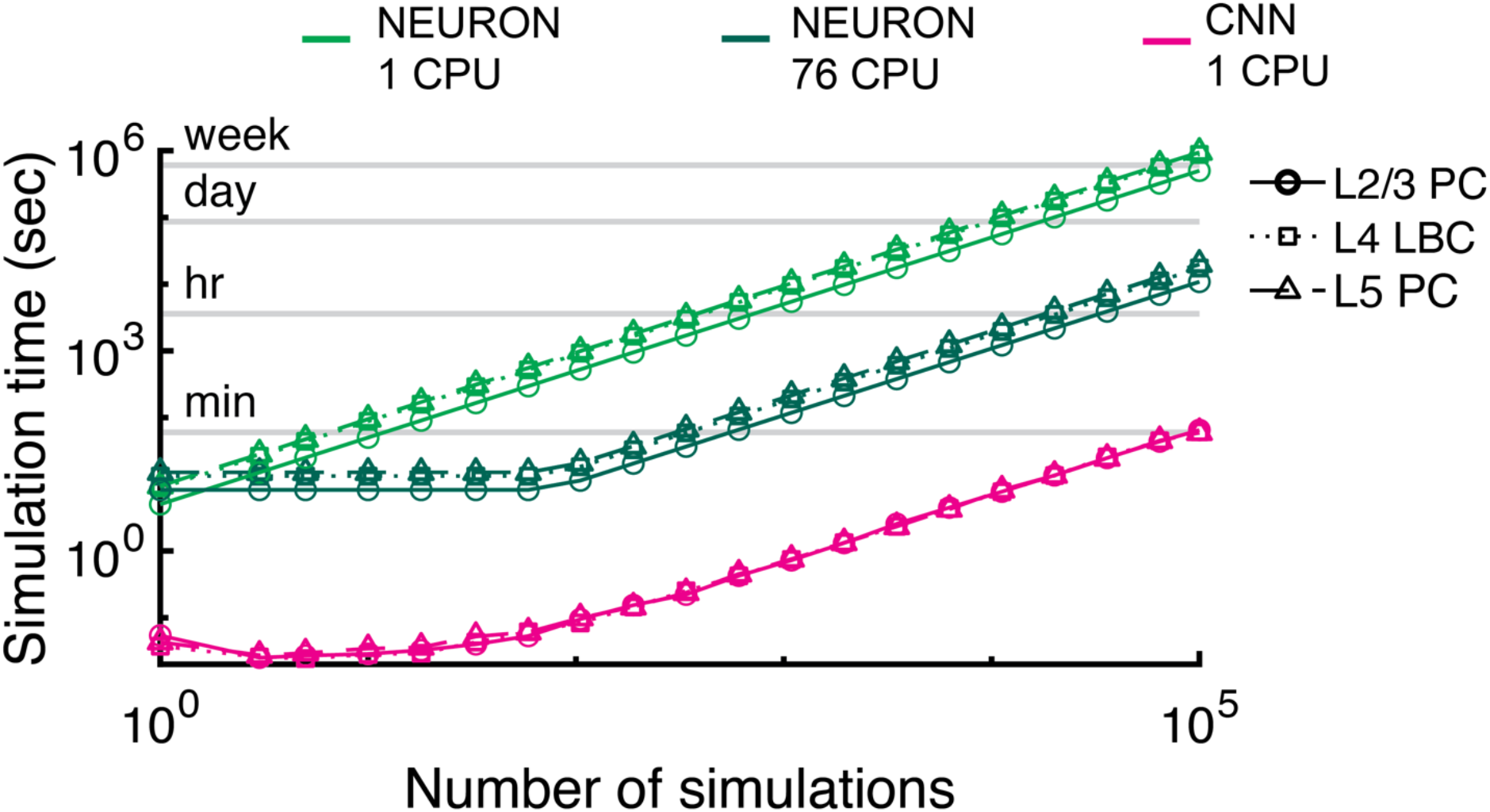
CNN threshold estimation is 2 – 4 orders of magnitude faster than NEURON. Simulation time for determining thresholds with binary search in NEURON on single CPU (light green), parallelized on single high-performance computing (HPC) node with 76 CPUs (dark green), or with CNN on single CPU (magenta) for example L2/3 PC (circles), L4 LBC (squares), and L5 PC (triangles). NEURON data points are based on average single threshold simulation time (*n* = 5) on a single CPU of laptop or HPC machine, and these were then extrapolated by assuming linear scaling (best case) and parallelization across all 76 CPUs for the latter approach. CNN data points are the actual run times for estimating the corresponding number of thresholds.

## 4. Discussion and conclusion

We developed a 3D CNN architecture that provided accurate estimates of the thresholds of biophysically realistic cortical neuron models for activation by TMS-induced E-fields. Using E-field vectors sampled on regularly spaced grids encompassing the neuronal morphologies, the CNNs predicted activation thresholds for TMS pulses with mean absolute percent error less than 2.5% for all models. The CNNs substantially outperformed an alternative approach in which the non-uniform E-field in the vicinity of the neurons was approximated as uniform, which yielded lower correlations and higher error. Reducing the number of E-field sampling points slightly increased error while still outperforming the uniform E-field method. The optimal E-field sampling grid size was determined by a balance between encompassing the spatial extent of the neuron without reducing significantly the sampling density or sampling extraneous distant points, e.g., E-fields outside the gray matter tissue volume. Representing the E-field vectors in spherical coordinates using four variables, compared to Cartesian coordinates using three variables, increased the error of the CNN predictions. Additionally, the CNNs were able to predict accurately thresholds for uniform E-fields, which were never seen during training, and the CNNs also performed well at predicting thresholds for point source stimulation, even though the CNNs were not trained on such data and the point source E-field is highly non-uniform compared to the TMS or uniform E-fields. On a single CPU, estimating thresholds with the CNN for 1000 unique E-field distributions, corresponding to different neuron locations and orientations in the brain, required only 670 ms, while the equivalent NEURON simulations would take, at minimum, 1.4 to 2.6 hours (depending on model complexity), providing three to four orders of magnitude speedup.

Our approach took advantage of the ability of CNNs to extract salient features from data with regular spatial structure, as exists in 2D and 3D images. The 3D grid of E-field vectors is analogous to a volumetric image with three color channels for each E-field component. By convolving the entire image with multiple local kernels, in this case 9 × 9 × 9 images convolved with 3 × 3 × 3 kernels, CNNs extract spatially invariant features of input images for processing by deeper layers with fewer parameters than fully connected 1D networks.

In the neuron simulations, action potentials were initiated at axon terminals aligned with the local E-field, and the estimated TMS thresholds were inversely correlated with the E-field magnitude, degree of branching, myelination, and diameter [19,49]. Therefore, for a given morphology, one would expect the exact location of the E-field vectors to be important for predicting the threshold, i.e., whether the E-field is high or low near an aligned axonal branch. The CNN predicted accurately thresholds for a wide range of non-uniform and uniform E-field inputs, with mean percent error close to the window used in the simulated binary search (2%), which suggests the CNN, to some degree, encoded features of the E-field distribution relative to an internal representation of the axonal geometry. CNNs are classically thought to be insensitive to absolute spatial location; however, several studies have shown CNNs can learn implicitly spatial information by exploiting differences in convolutional kernel activations near image borders [50–52].

Several factors may have contributed to the residual error and outliers. One factor was likely the under-sampling of E-field variations in regions with higher gradients or near tissue boundaries (i.e., gray matter / white matter and gray matter / CSF), as errors were increased by reducing the sampling resolution while keeping the CNN architecture fixed. Using fewer sampling points would be advantageous to reduce the time spent interpolating E-field vectors within the FEM mesh, which can be substantial for high sampling densities and large neuronal populations. For reference, the NEURON simulations require interpolating E-field vectors at all compartments, which number over 880 for all cell types, except one L2/3 PC with 610 compartments, and are as high as 4,196 for one of the L4 LBCs, due to its dense axonal arborization. Still, even the 9 × 9 × 9 CNN E-field sampling grid (729 E-field vectors total) required fewer E-field samples than most of the biophysically realistic neuron simulations. Reducing the sampling resolution of the E-field distributions provided the network with less accurate representations of the E-field gradients and the E-field strength and direction at the site of AP initiation. Metrics related to these three factors explained nearly 30% of the variance in CNN prediction error at the lowest sampling resolution. The CNNs were able to generalize well to predict thresholds for uniform E-field (zero spatial gradient), but, as expected, prediction errors were significantly higher for the point source E-fields, which had over 100 times higher spatial gradients. Nonetheless, the CNNs reproduced qualitatively the current–distance relationship without retraining, suggesting the models could learn to estimate more accurately responses to intracortical microstimulation, as well. This would require training a set of CNNs using the desired pulse waveform (e.g., 0.2 ms cathodic square pulse) and potentially using a higher E-field sampling resolution, although alternative approaches to spatial and temporal representation of the E-field can also be considered, as discussed below.

TMS thresholds are strongly correlated with the E-field magnitude [14,18,53], so we expected that converting the E-field to spherical coordinates would reduce error further, since the magnitude is explicitly represented as one of the coordinate variables. However, the CNNs performed best with Cartesian E-field coordinates. This was possibly due to the additional E-field coordinate variable introduced to represent the azimuthal component; while avoiding circular discontinuities in the data [54,55], the use of spherical coordinates increased the dimensionality of the data without providing more information to the network [56]. Furthermore, prediction errors were slightly higher in the L2/3 PCs compared to the L4 LBCs and L5 PCs. The L2/3 PCs not only had a wider range of E-field thresholds due to their high threshold anisotropy (i.e., strong dependence on orientation with respect to the E-field), but they also have long horizontal axon branches that extend several millimeters at oblique and tangential angles. This made it difficult to select a grid size that encompassed the neuron models without also sampling from points in the CSF tissue volume of the FEM, potentially resulting in some positions where the CNN did not have adequate E-field samples near the activated axonal branch. Using the regular sampling grid allowed the same grid points to be used for every clone at each position and reduced the number of E-field points to interpolate by a factor of five. However, one limitation of this approach is the sampling grid shape does not always conform to the anatomy of the neuron or the cortex, depending on neuron depth, local curvature, and thickness of the cortical sheet. An alternative approach might be to instead represent the E-field on the neural compartments, e.g. using graph convolutional neural networks [57], as the model morphologies are more reliably positioned within the gray matter. Sampling along the morphology may also more efficiently capture variations of the E-field magnitude at relevant locations without including irrelevant E-field vectors distant from an axonal branch or E-field vectors that fall in a different tissue compartment. This approach, however, will require more preprocessing of the E-field simulations to obtain unique spatial sampling for each cell morphology and orientation.

This is the first study to use ANNs to represent the response of morphologically realistic, multicompartmental neuron models to stimulation with exogenous E-fields. Previously, Chaturvedi et al. used an ANN with one hidden layer to estimate the volume of tissue activated (VTA) by multi-contact configurations of deep brain stimulation as computed with multicompartmental straight axon models [58]. Otherwise, recent studies used deep learning to predict the subthreshold and suprathreshold temporal dynamics of neuron models in response to synaptic inputs [27,28]. Olah et al. tested the ability of multiple ANN architectures to predict somatic voltage and current time series of biophysically realistic models of L5 PCs in NEURON, similar to those included in this study [27]. They found the only architecture capable of predicting accurately the subthreshold and suprathreshold dynamics was a model with both convolutional and long short-term memory (CNN-LSTM). Interestingly, the CNN-LSTM reliably reproduced the response of the NEURON models to distributed synaptic inputs using only the somatic voltage. At the same time, it is unclear whether this approach could predict efficiently the dynamics in the rest of the neuronal compartments, which is necessary for applications involving extracellular stimulation, as in the present study. Beniaguev et al. conducted a similar study and found a temporal convolutional network (TCN) could reproduce the input–output properties of a biophysically detailed L5 PC [28]. Our focus on estimating thresholds for activation allowed a simpler implementation that resulted in low prediction error, indicating the CNNs learned the dependency of activation thresholds on E-field spatial distributions without requiring explicit representation of underlying neural dynamics, i.e., voltage and current time courses in any compartment. Still, these studies suggest ANNs can accurately represent the spatiotemporal computations performed by spatially extended neuron models. In our approach, pulse waveform was implicit in the training data, and this requires separate CNNs to be trained on threshold data with other pulse waveforms. For most TMS applications, this approach is likely acceptable, due to the limited range of pulse waveforms produced by conventional TMS devices (e.g., monophasic and biphasic pulses). However, a time-dependent approach is necessary to estimate thresholds for novel pulse waveforms, such as those generated by the specific inductance and resistance of novel coils or by TMS devices with controllable pulse shape [59,60], without requiring separate training sets and CNNs for discrete waveform selections. Additionally, the calculation of “ground truth” thresholds was conducted with quiescent neurons, and endogenous activity can alter the response to TMS experimentally and shift the thresholds of individual neurons [61– 63]. Combining 3D convolutional layers to encode the interaction between the extracellular E-field and the neuron morphology with recurrent layers to capture temporal dynamics of the pulse and/or the neural membrane could combine the advantages of both approaches and is likely an important direction for future research.

The CNNs provided several orders of magnitude reductions in required computation, making it feasible to incorporate neural response models in E-field simulation packages for use on commonly available computational resources. Simulating the TMS-induced E-field in head models with 1^st^ order FEM, as used in SimNIBS [29], requires on the order of 1–2 minutes on a typical computer [64]. Estimating the response of the full population we modeled in the *ernie* mesh (3 layers, 5000 neurons per layer, 12 rotations) would add less than 3 minutes on a single CPU. This time would be dramatically reduced by using a GPU, but even if a GPU is not available, further speedup on CPUs is feasible by optimization of the CNN implementation.

In conclusion, subject-specific head models of the E-field can support accurate dosing and targeting of cortical structures by TMS; however, these models alone cannot predict the physiological response. Combining E-field models with biophysically realistic neuron models addresses this limitation, but calculating the neural response is extremely computationally demanding [18]. Combining ANN estimators of the neural response with fast approaches to E-field computation [24,65,66] may enable more TMS users to adopt these multi-scale biophysically-based models in research and clinical applications.

## 5. Acknowledgements

This work was supported by NIH grants R01NS088674, R01NS117405, and R01MH128422, and NSF Graduate Research Fellowship DGF 1106401. The content is solely the responsibility of the authors and does not necessarily represent the official views of the funding agencies. We thank Dr.

Boshuo Wang, Dr. Nicole Pelot, Minhaj Hussein, and Dr. David Carlson, for their technical assistance and helpful discussions. Dr. Wang also assisted in revising early drafts of the manuscript. We also thank the Duke Compute Cluster team for computational support.

## 7. Supplementary Figures

**Supplementary Figure S1.**
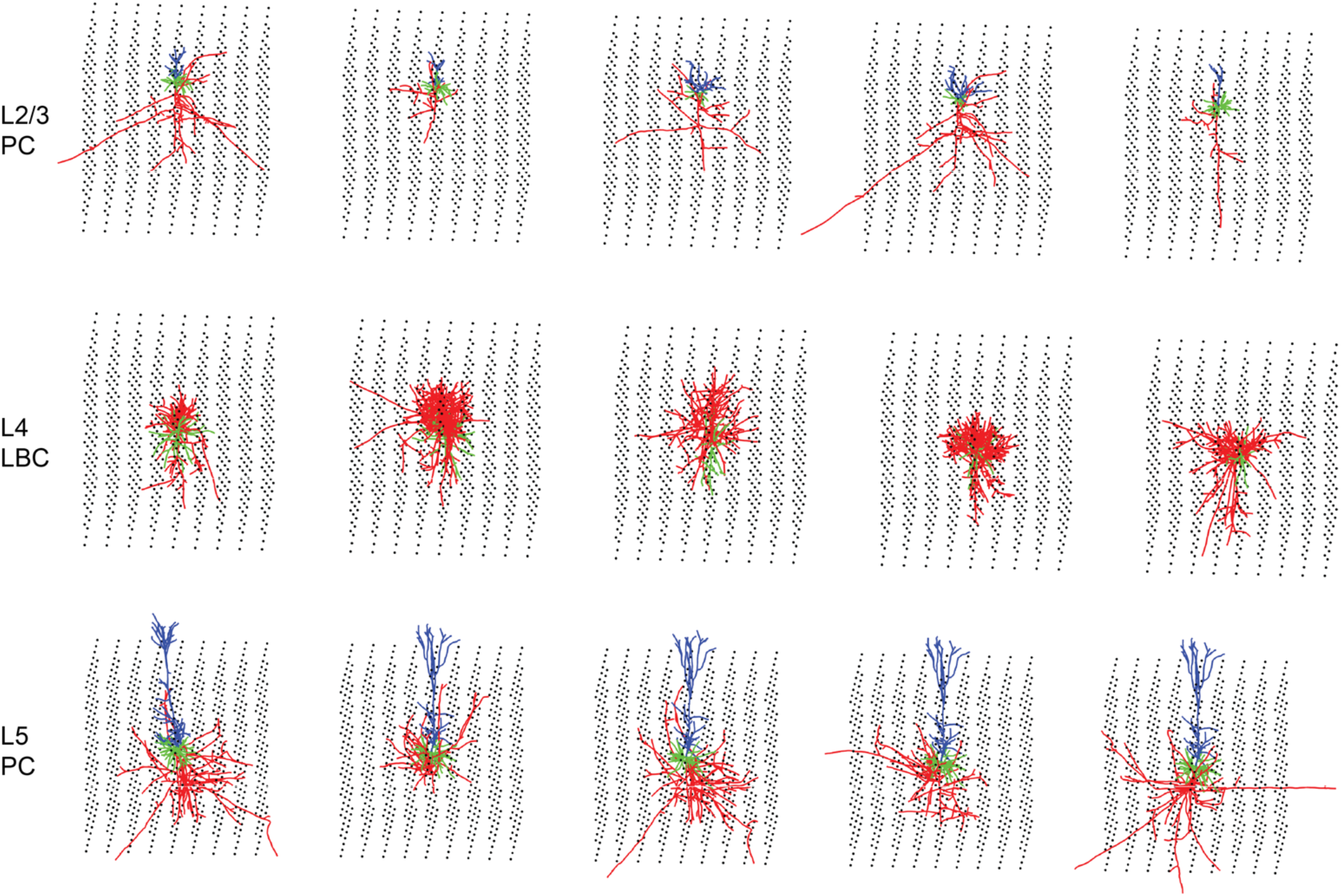
E-field sampling grids for all model neurons. E-field sampling grid for **A)** L2/3 PCs with side length *l* = 2 mm shifted in z-direction by −0.49 mm; **B)** L4 LBCs with side length *l* = 1.5 mm; and **C)** L5 PCs with side length *l* = 1.5 mm. All grids shown have *N* = 9 points per dimension.

**Supplementary Figure S2.**
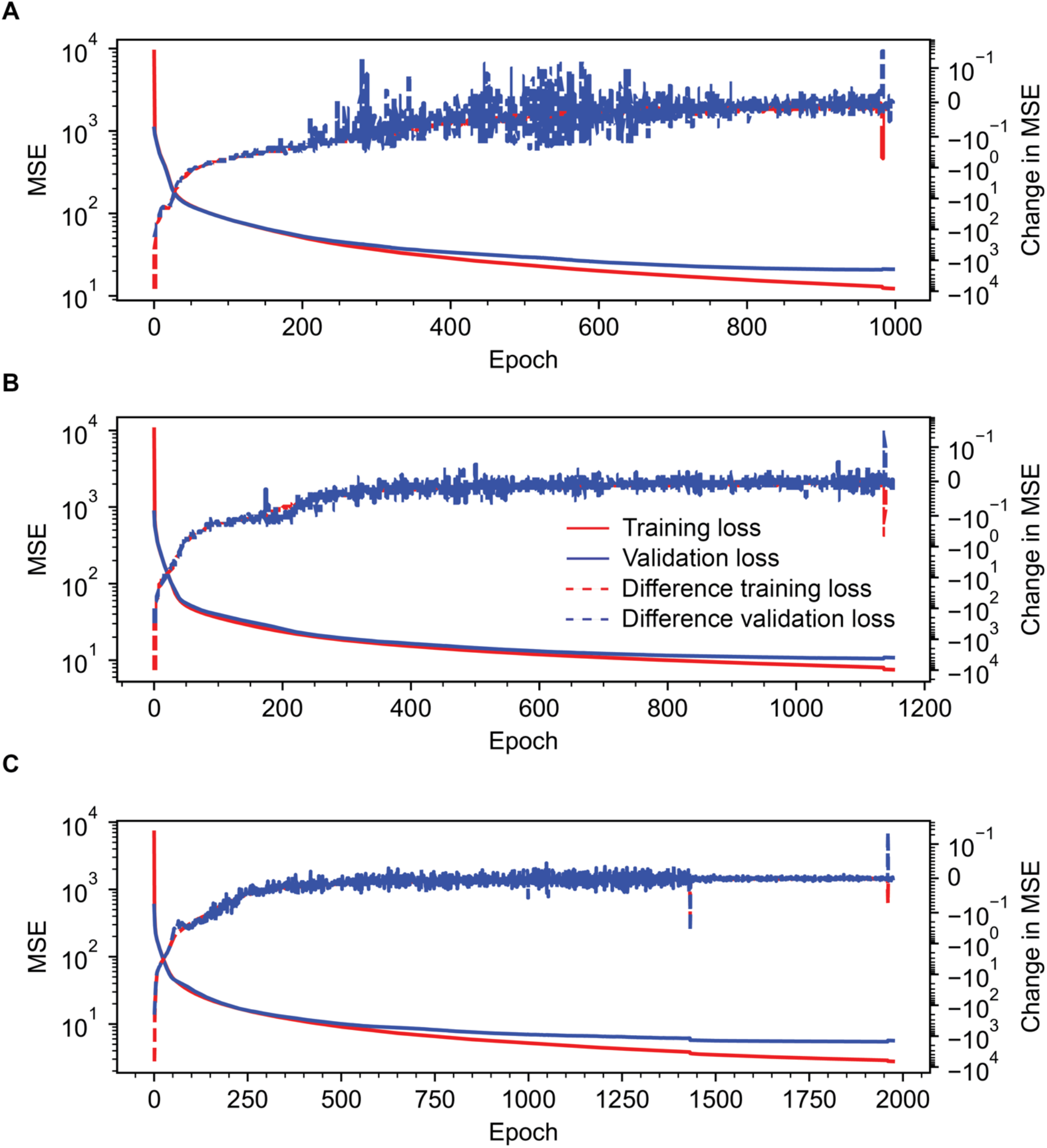
Example CNN training curves. Training and validation loss, quantified as mean squared error (MSE), change in loss per training epoch for **A)** L2/3 PC (clone 1), **B)** L4 LBC (clone 1), and **C)** L5 PC (clone 1). Change in loss is plotted on symmetrical log scale with linear range between ± 10^−1^. For reference to corresponding clone’s morphology, see Supplementary Figure S1.

**Supplementary Figure S3.**
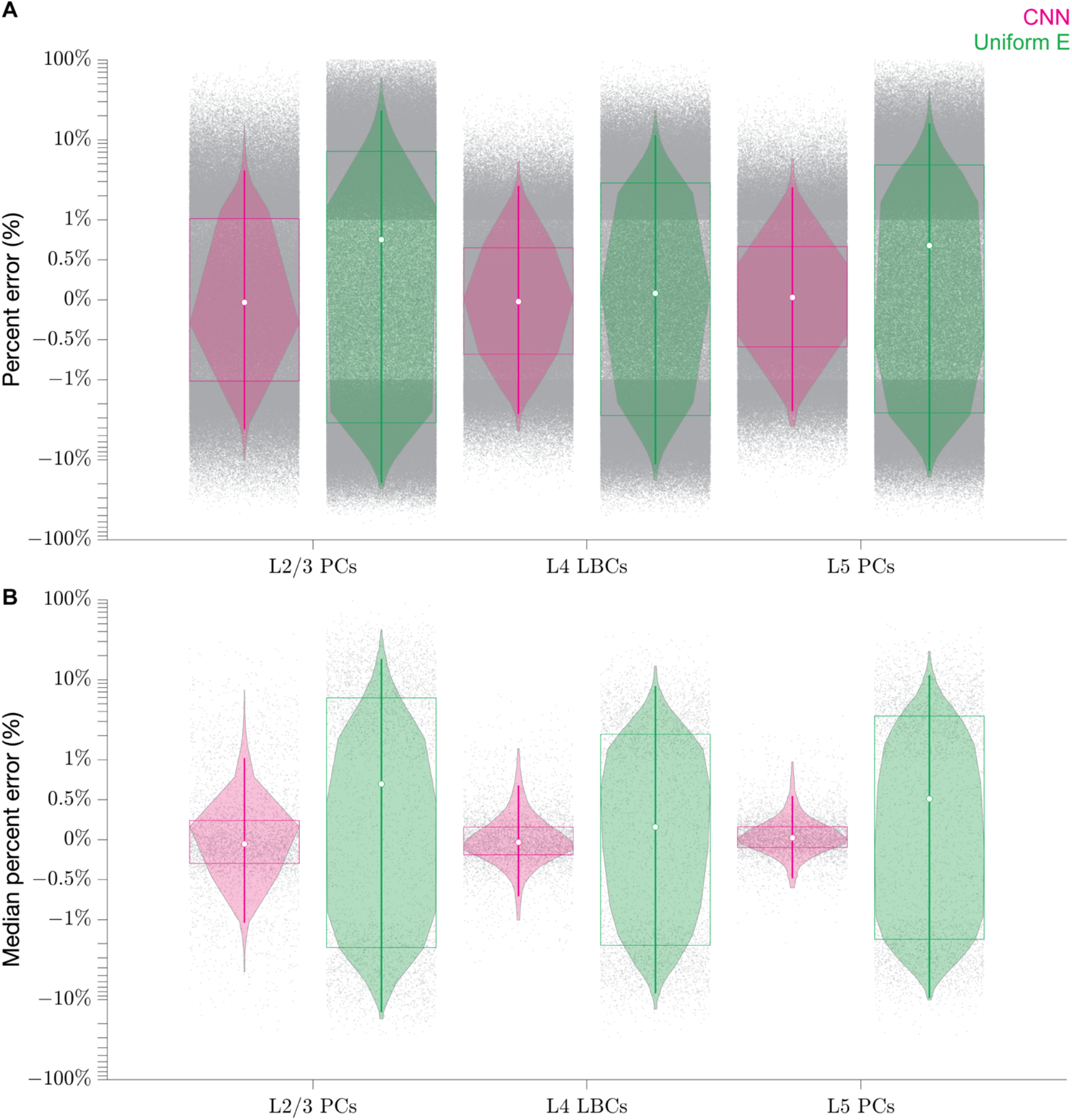
Distribution of test prediction error for CNN and uniform E-field method. Distributions of **A)** percent errors of all clones and rotations and **B)** median percent errors across clones and rotations at each position shown with data points (gray); estimated probability kernel density (violin plots) spanning 98% of data; and box and whisker plots indicating median (white point), 1^st^ and 3^rd^ quartiles (rectangular box), and whiskers (vertical lines) extending to 1.5 × interquartile range below and above 1^st^ and 3^rd^ quartile, respectively. Note the log-linear-log vertical axis with linear scaling between -1 and 1% and logarithmic scaling outside this range.

**Supplementary Figure S4.**
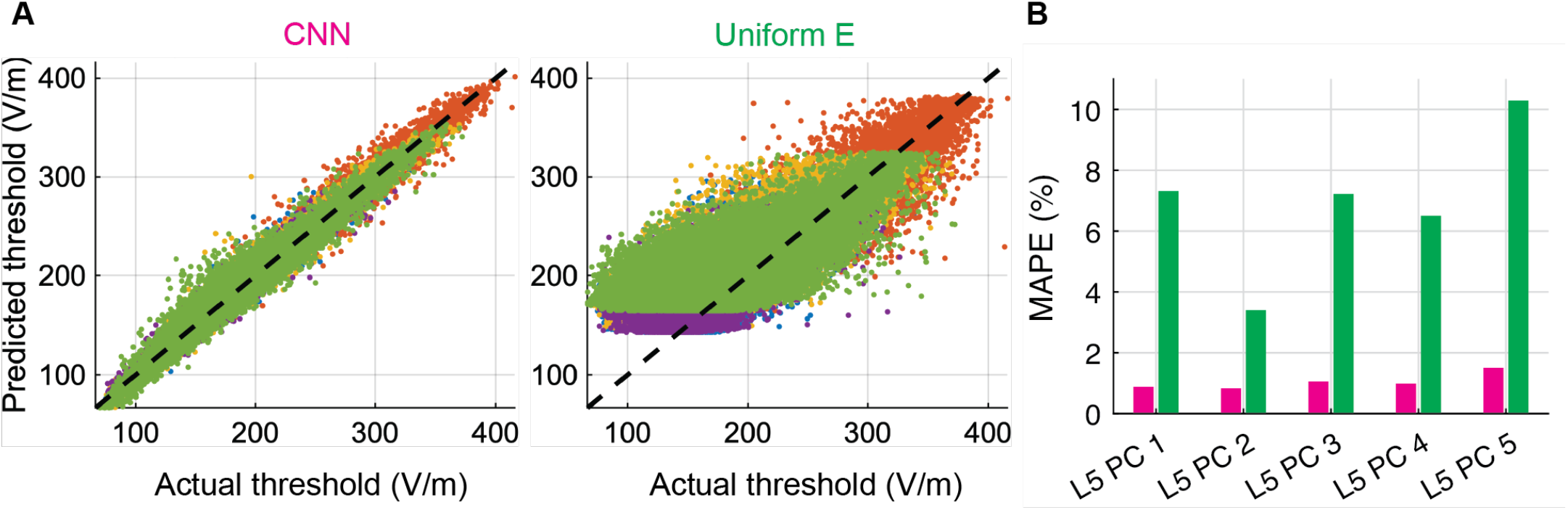
CNN also predicts accurately thresholds for biphasic TMS pulse. **A)** Predicted threshold E-field at soma of all five L5 PC clones for MagProX100 biphasic TMS pulse by CNN (left column) and uniform E-field approximation (middle column) across entire test dataset plotted against NEURON simulation thresholds (actual). **B)** Mean absolute percent error (MAPE) on test dataset for CNN (magenta) and uniform E-field approach (green), separated by clone.

**Supplementary Figure S5.**
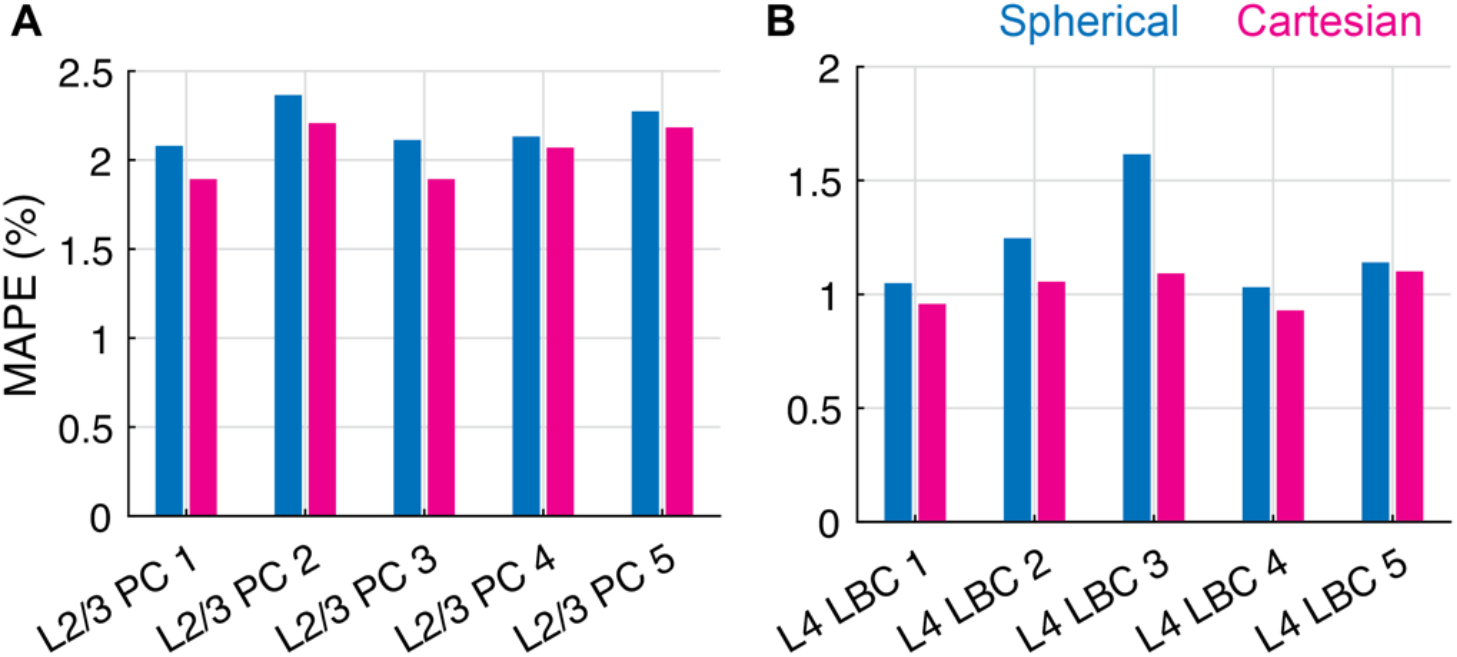
Effect of E-field vector coordinate system on performance for L2/3 PCs and L4 LBCs. Mean absolute percent error (MAPE) metric on test dataset for **(A)** L2/3 PC and **(B)** L4 LBC CNNs with E-field input represented with either spherical coordinates (pre-processing described in Section 2.2.1) or Cartesian coordinates.

**Supplementary Figure S6.**
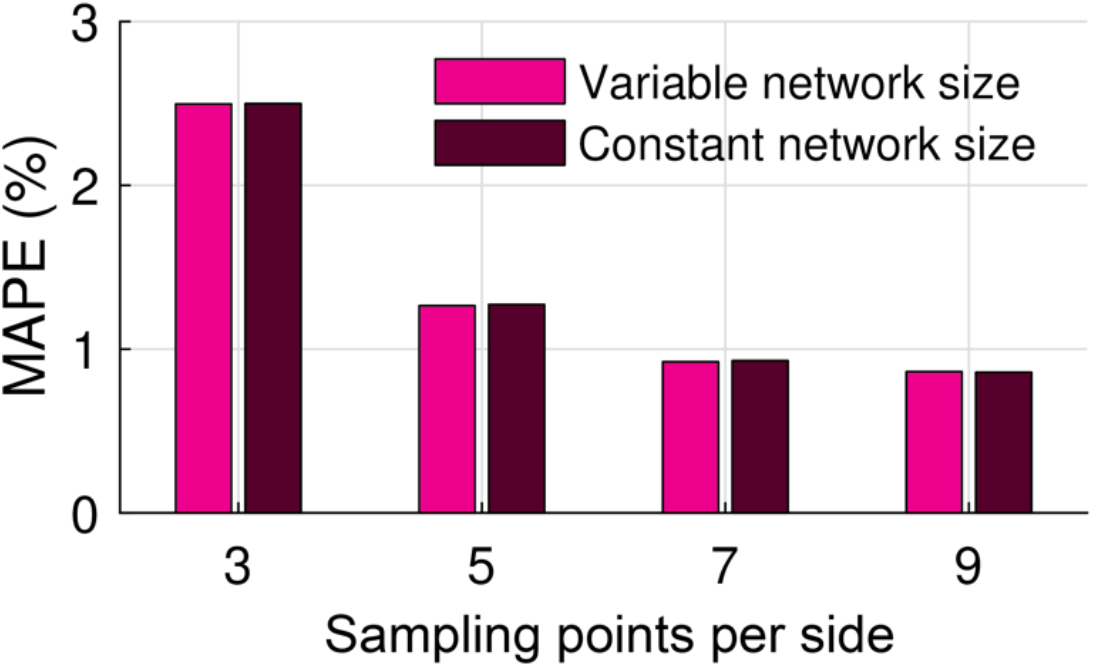
CNN error increases with fewer E-field sampling points for both variable and constant network size. MAPE metric for example L5 PC (clone 1) CNN for different sampling resolutions using either variable network size, in which the architecture is modified to accommodate lower resolution inputs, or constant network size (see Section 2.2.3), in which the architecture is kept constant and the inputs are upsampled to the highest resolution (*N* = 9).

**Supplementary Figure S7.**
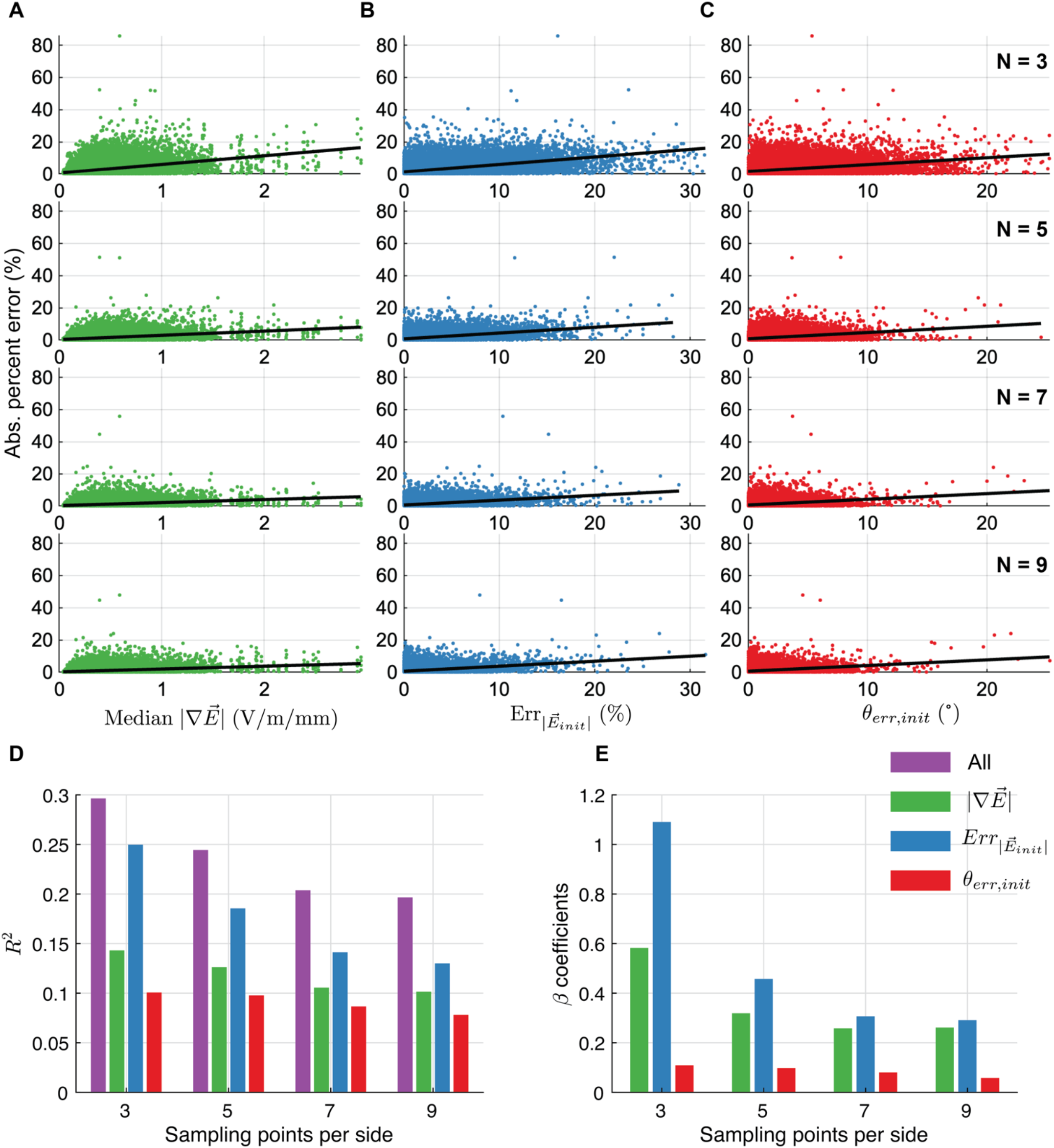
CNN error correlates with E-field gradient and E-field at AP initiation site. CNN prediction errors for test dataset of example L5 PC (clone 1) plotted against **A)** median magnitude of directional E-field gradient, **B)** absolute percent error of E-field magnitude at AP initiation site, and **C)** error of E-field direction at AP initiation site for *N* = 3, 5, 7, or 9 sampling points per dimension, with single regression lines overlaid. Median E-field gradients in A) were all calculated using E-field grids with *N* = 13 sampling points per dimension. For definition of E-field magnitude and direction error metrics in B) and C), see section 2.2.1. **D)** *R*^2^ values for multiple linear regression with all metrics (purple) and single regression with the metrics in A–C for each sampling resolution. Adjusted *R*^2^ used for the multiple linear regressions to account for the effect of adding model predictors. **E)** Standardized *β* coefficients for multiple linear regression with all metrics in A–C at each sampling resolution.

**Supplementary Figure S8.**
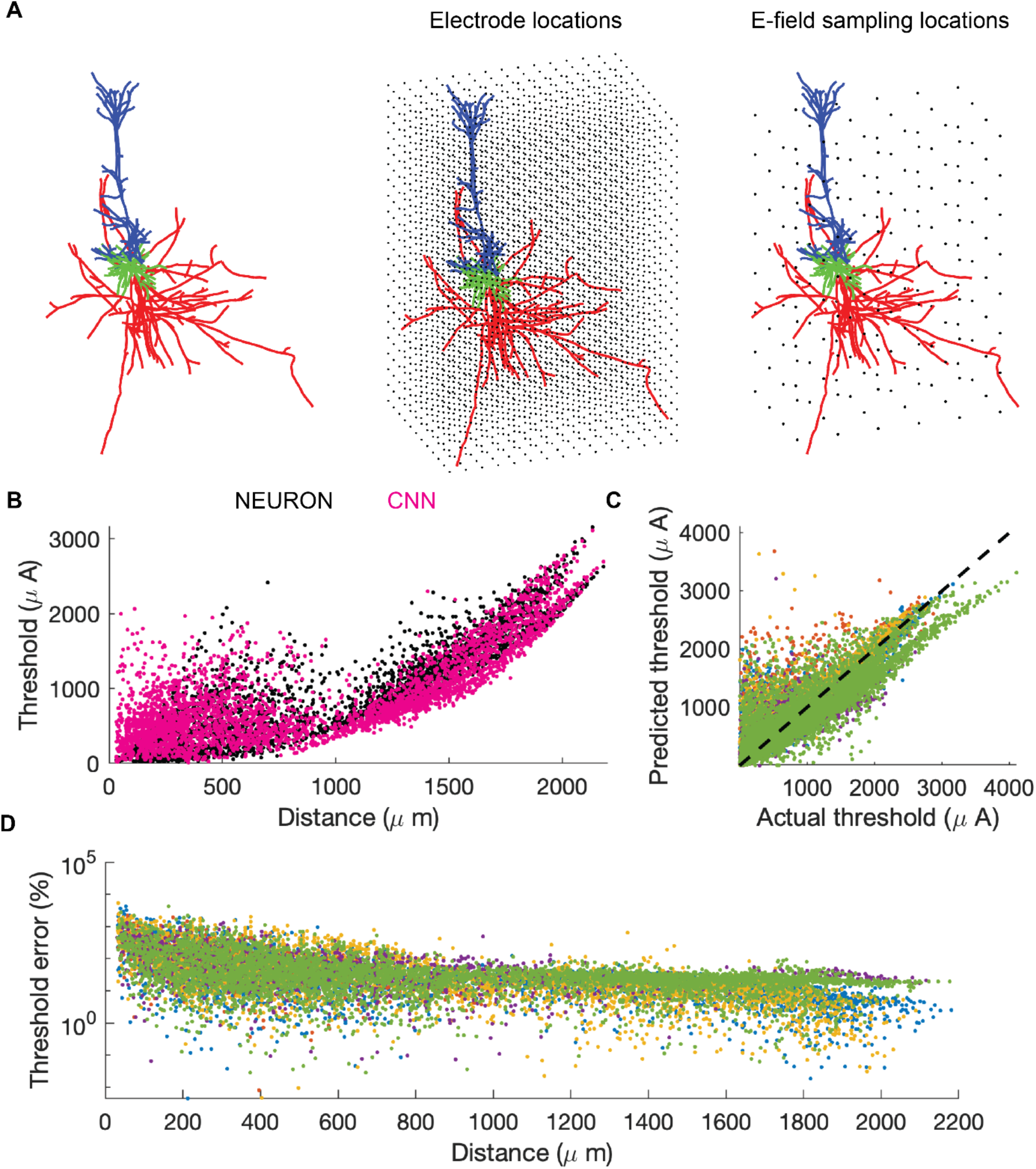
CNN trained on TMS-induced E-field predicts response to point source with reduced accuracy. We modeled stimulation with a microelectrode as a point current source in a homogenous, isotropic medium with conductivity *σ* = 0.276 S/m, as in [19], and computed thresholds in NEURON for electrode locations throughout a 3D grid encompassing each of the L5 PC morphologies in 100 µm steps. Electrode locations within 30 µm of a neuronal compartment and eliciting dendritic activation at lowest threshold were excluded. We used the same MagProX100 monophasic TMS pulse to match the pulse waveform used with the CNNs. We estimated thresholds with the CNNs pretrained on TMS thresholds by inputting the E-field distribution generated for each electrode location, normalized to the magnitude of the center grid point. The E-field per unit current is given by 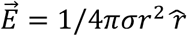, where *r* is the electrode-to-sampling-point distance and 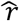 is the unit vector in the radial direction in spherical coordinates. **A)** Example L5 PC morphology (left) with electrode locations overlaid (middle) or E-field sampling points for CNN (right). **B)** Threshold current–distance plot for example L5 PC (clone 1) generated with NEURON simulations or the trained CNN. Each point represents the threshold for a different electrode location within the 3D grid and the distance from that electrode to the point of action potential initiation. **C)** Predicted current thresholds by CNN plotted against NEURON simulation thresholds (actual) for all five L5 PC clones. The correlations were weaker than for the TMS simulations, but were still significant, with *R*^2^ ranging from 0.617 – 0.792 (*p* < 0.001). **D)** Threshold percent error plotted against distance for all five L5 PC clones, demonstrating lower errors for more distant electrode locations. This was likely due to the CNNs being trained on E-fields with low spatial gradients and the decrease in spatial gradient with distance from a point source.

## Footnotes

1 The estimate of 154 million neurons was calculated based on a neocortical neuron density of 120,000 per mm^2^ (through the cortical depth) [67] and the surface area of the precentral gyrus of 1,280 mm^2^ in the SimNIBS v1.0 *almi5* example head mesh [29].

2 Single CPU run times in NEURON varied between the different CPUs tested, with faster serial run times on the Macbook Pro (shown in Figure 7). On the HPC node, single CPU run times were 6.6 ± 0.3 sec for the L2/3 PC, 10.5 ± 0.3 sec for the L4 LBC, and 11.2 ± 0.2 sec for the L5 PC.

